# LLMsFold: Integrating Large Language Models and Biophysical Simulations for De Novo Drug Design

**DOI:** 10.64898/2026.03.02.709055

**Authors:** Wageesha Widuranga, Davide Rigoni, Fabio Bove, Dario Righelli, Salvatore Romano, Rosa Visone, Marilena V. Iorio, Marco Grassia, Giuseppe Mangioni, Pietro Liò, Cristian Taccioli

## Abstract

The discovery of novel small molecules is challenging because of the vastness of chemical space and the complexity of protein-ligand interactions, leading to low success rates and time-consuming workflows. Here, we present LLMsFold, a computational framework that combines Large Language Models (LLMs) and biophysical foundation tools to design and validate new small molecules targeting pathogenic proteins. The pipeline starts by identifying viable binding pockets on a target protein through geometry-based pocket detection. A 70-billion-parameter transformer model from the Llama family then generates candidate molecules as SMILES strings under prompt constraints that enforce drug-likeness. Each molecule is evaluated by Boltz-2, a diffusion-based model for protein-ligand co-folding that predicts bound 3D structure and binding affinity. Promising candidates are iteratively optimized through a feedback loop that prioritizes high predicted affinity and synthetic accessibility. We demonstrate the approach on two challenging targets: ACVR1 (Activin A Receptor Type 1), implicated in fibrodysplasia ossificans progressiva (FOP), and CD19, a surface antigen expressed on most B-cell lymphoma and leukemia cells. Top candidates show strong in silico binding predictions and favorable drug-like profiles. All code and models are made available to support reproducibility and further development.

## 1 Introduction

Drug discovery remains a challenging and time-consuming endeavor. Although the synthetically accessible chemical space is vast, only a small fraction of compounds typically meets the criteria for a successful drug Bohacek et al. [1996], Polishchuk et al. [2013], Walters and Murcko [2002]. Traditional de novo design methods have struggled with this challenge because early algorithms produce candidates with poor pharmacokinetics or synthetic feasibility [Schneider and Fechner, 2005]. Ligand-based virtual libraries combined with heuristic scoring have brought improvements [Hartenfeller and Schneider, 2010], but remain limited by rigid rules and the need for expert curation. In recent years, deep generative models have demonstrated the ability to generate novel chemical structures more quickly and efficiently. Notably, transformer algorithms [Vaswani et al., 2023], the same architectures that revolutionized natural language processing, can be used to generate chemical representations as a language, learning the grammar of chemical structures. This has paved the way in drug design [Zhavoronkov et al., 2019]: models such as DrugGPT (a GPT-based ligand generator) [Li et al., 2023] have demonstrated the ability to propose new and synthetically accessible molecules predicted to bind specific protein targets [Liu et al., 2023, Grisoni et al., 2021]. By training on vast datasets of known drugs and bioactive compounds, these large language models might capture structural patterns and activity relationships, enabling them to generate drug-like SMILES [Weininger, 1988, Weininger et al., 1989, Weininger, 1990] strings in a shorter time. LLMs’ Despite these advances, using LLMs for drug design presents key challenges. The models must respect chemical validity and generate compounds with realistic chemical properties for a quick and efficient synthesis. Moreover, predicting true binding affinity usually requires physics evaluation beyond the LLMs capabilities. In particular, the Boltz family of models, developed by MIT & Recursion, introduced a new model by predicting full protein complexes with accuracy approaching that of AlphaFold [Abramson et al., 2024]. The first model, Boltz-1 [Wohlwend et al., 2024], achieved AlphaFold3-level performance on diverse protein/protein and protein/ligand benchmarks and was released open-source. Its successor, Boltz-2 [Passaro et al., 2025], further integrates geometric deep learning with diffusion-based prediction algorithms to handle all-atom interactions in protein complexes. It is able to predict binding pockets, providing confidence metrics (pLDDT, interface TM-score, etc.), making it a powerful tool that is an alternative to traditional docking [Trott and Olson, 2010]. In particular, Boltz-2 not only predicts how a ligand binds, but it also provides an affinity estimate and effective virtual screening at speeds that exceed classical simulations [Cournia et al., 2017, Schindler et al., 2020].

In this work, we combine these innovations, that is, LLM-based molecular generation and Boltz-2 affinity evaluation, into an integrated pipeline, which we call LLMsFold. In particular, rather than training a model from scratch for a specific task, we provide the language model with a few examples of clinically relevant molecules, including those in clinical trials, approved drugs, or well-studied compounds, directly in the prompt. The model then learns from these examples what it needs to achieve.

In the literature, several approaches exist for generating small molecules conditioned on a target pocket. State-of-the-art methods include Pocket2Mol Peng et al. [2022], REINVENT 4 Blaschke et al. [2020], Loeffler et al. [2024], and SynFlowNet Cretu et al. [2025], which explore different paradigms for target-guided molecular design. Pocket2Mol operates directly in 3D pocket space, REINVENT 4 generates SMILES strings through reinforcement learning, and SynFlowNet is based on GFlowNets Bengio et al. [2023], whose action space uses chemical reactions and purchasable reactants to sequentially build new molecules. While these methods have shown strong performance, they often require target-specific adaptation and considerable computational cost compared to our approach

In this study, we focus on two biomedically relevant targets: ACVR1, a kinase whose gain-of-function mutation (mutated in R206H) drives a rare genetic disease (FOP or Fibrodysplasia Ossificans Progressiva) [Shore et al., 2006], and CD19 [Tedder and Isaacs, 1989], a B-cell surface antigen that regulates cell proliferation in leukaemia [Awan et al., 2010] and lymphoma [Zhang et al., 2019], for which new small-molecule modulators could be a valid alternative to recent therapies [June and Sadelain, 2018]. These diseases respond to established pharmacological treatments, albeit not curatively, and therefore represent a suitable benchmark for de novo drug design. We demonstrate that LLMsFold can generate molecules that pass successive filters: (1) they fit sterically into a predicted binding pocket on the target protein; (2) they are chemically drug-like; (3) they bind the target with high predicted affinity and specific pose; and (4) they are synthetically accessible and novel. Iterative optimization using a feedback loop from Boltz-2 and cheminformatics scoring further refines the candidates toward optimal predicted inhibitors.

By combining LLM-based generation and Boltz-2 affinity evaluation, our approach aims to address issues of classical drug design and discovery. In the following sections, we detail the LLMsFold methodology and present results highlighting several lead compounds for ACVR1 and CD19 pockets. We also discuss the broader implications of this AI-pharmacology hybrid strategy for accelerating early-stage drug discovery, where the ability to rapidly explore and validate chemical space could significantly improve efficiency and reduce the timeline for discovering new therapeutics.

## 2 Materials and Methods

This section describes the complete *LLMsFold* framework, which is visible in Figure 1. The pipeline integrates structure-based binding pocket identification, large language model (LLM)-guided molecular generation, biophysical evaluation through Boltz-2 co-folding simulations, iterative feedback-driven optimization, and cheminformatics validation. Starting from experimentally resolved protein structures obtained from the Protein Data Bank (PDB), the workflow automatically identifies candidate binding regions, generates novel SMILES-based molecular structures conditioned on pocket characteristics, and evaluates their predicted binding affinity and structural plausibility. Iterative optimization further refines candidate molecules using feedback derived from affinity prediction and molecular similarity metrics. Finally, generated compounds are filtered using standard drug-likeness, synthetic accessibility, and novelty assessment criteria to prioritize chemically viable lead candidates.

**Figure 1:**
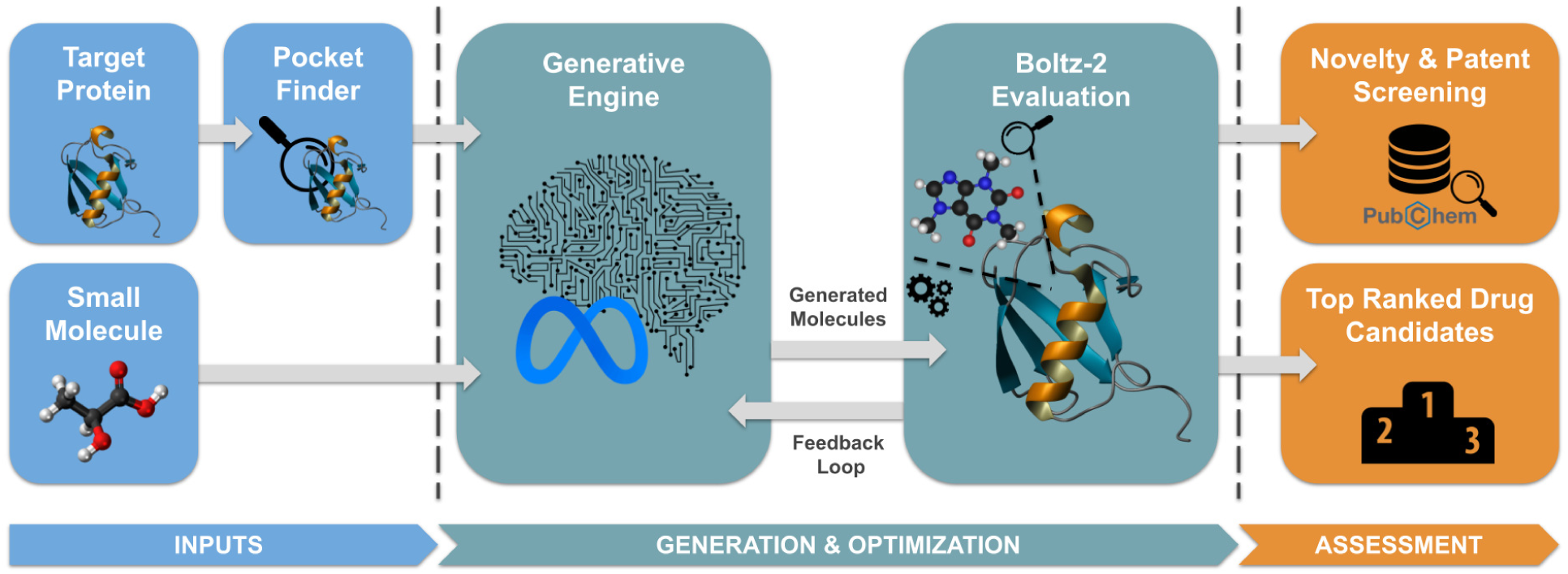
The LLMsFold Framework. An integrated lead discovery pipeline transitioning from geometric pocket identification to LLaMA-driven molecular generation. The workflow integrates Boltz-2 co-folding for 3D structural validation and binding affinity prediction, culminating in an iterative feedback loop for multi-parameter optimization of drug-like candidates.

### 2.1 Binding Pocket Identification

The LLMsFold pipeline begins with the identification of potential binding sites, requiring only the structure of the target protein in PDB (Protein Data Bank) format sourced from RCSB PDB [Berman et al., 2000]. Rather than assuming a predefined active site, we employ the Convex Hull Pocket Finder [Saberi Fathi and Tuszynski, 2014], implemented within the DeepChem library [Ramsundar et al., 2017], to scan the molecular surface for concave regions capable of accommodating ligands. This algorithm systematically identifies potential cavities based strictly on spatial coordinates. In particular, the algorithm identifies clusters of surface points representing potential binding pockets. For each identified region, we define a three-dimensional bounding box that minimally encloses it. The parameters are calculated as: (i) the midpoint of the extremal coordinates, and (ii) the side lengths representing the spatial span in each dimension. These parameters are exported in a standardized CSV format to ensure reproducibility of the search space in subsequent docking and simulation steps.

To prioritize drug-like binding sites, we apply a deterministic multi-stage filtering procedure. First, cavities with any box dimension smaller than 8 Å are excluded, ensuring that the pocket is sufficiently large to accommodate a typical drug-like molecule (*∼* 500 Da). Among the remaining candidates, we select the pocket with the minimum volume that still satisfies the dimensional constraints. This strategy minimizes the computational search space while preserving biologically relevant binding regions.

The selected binding pocket is subsequently refined to better account for protein flexibility during docking simulations. Specifically, the bounding box is isotropically expanded by 5 Å in all directions to accommodate potential side-chain movements, while the total box size is capped at 30 Å to maintain computational efficiency. We then identify residues constituting the chemical environment of the pocket using two complementary criteria: (i) residues containing at least one heavy atom within the expanded box, and (ii) residues with at least one atom located within 8 Å of the pocket center. Finally, the pocket center coordinates and box dimensions are provided as input to the docking simulation (see Table 1).

**Table 1:**
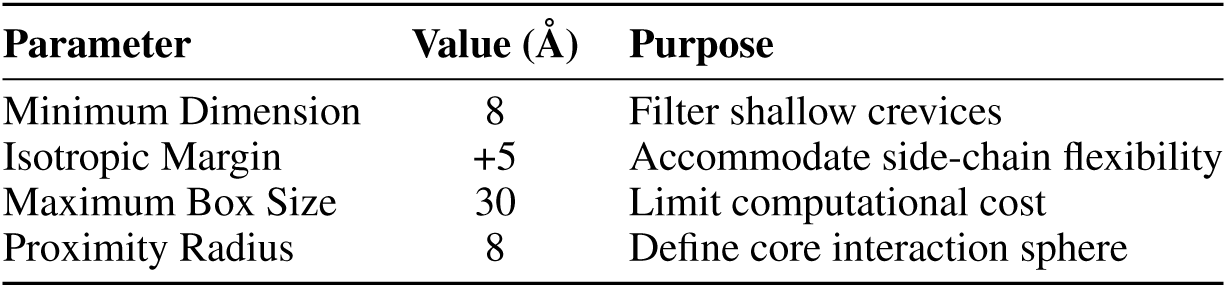
Default Geometric Parameters for Pocket Identification

| Parameter | Value (Å) | Purpose |
| --- | --- | --- |
| Minimum Dimension | 8 | Filter shallow crevices |
| Isotropic Margin | +5 | Accommodate side-chain flexibility |
| Maximum Box Size | 30 | Limit computational cost |
| Proximity Radius | 8 | Define core interaction sphere |

### 2.2 Generative Molecule Design via LLM (In-Context Learning)

Once we have identified the binding pocket on the target protein, we shift to generating the discovery of novel molecules that could bind to that particular pocket. For this, we use Llama-3.3-70B (Groq endpoint identifier llama-3.3-70b-versatile), a 70-billion-parameter language model from Meta AI. We chose this version because larger models are better at maintaining chemical validity across entire molecules. The algorithm treats molecule design as a language problem. Molecules are represented as SMILES [Weininger, 1988, Weininger et al., 1989, Weininger, 1990] strings, where atoms and bonds are encoded as characters, e.g., aspirin is represented as CC(=O)Oc1ccccc1C(=O)O. SMILES notation also captures structural features such as ring closures, branching patterns, aromatic systems, and bond connectivity through a compact sequence of symbols and brackets [Kaplan et al., 2020, Wei et al., 2022]. During its extensive training on diverse text, Llama has implicitly learned chemical patterns in which atoms commonly bond together, how rings are formed, and what functional groups frequently occur in chemical compounds [Brown et al., 2020]. This allows the model to generate new SMILES that represent chemically valid molecules, similar to how it works in natural languages.

Instead of retraining or fine-tuning the model, we use in-context learning [Dong et al., 2024]. This means we provide the model with a carefully designed prompt that includes a few examples of what we want, i.e., small molecules already used in clinical practice. The prompt begins with instructions describing the task and emphasizing key medicinal chemistry principles, particularly Lipinski’s Rule of Five for oral drug-likeness [Lipinski et al., 2001, Lipinski, 2004], and the need to avoid problematic substructures known as PAINS (Pan-Assay Interference Compounds) that cause false positives in biological assays [Baell and Holloway, 2010].

Regarding the example molecules provided to the model, we supply a single pocket-residue description shared across the whole prompt, together with a flat list of reference inhibitor SMILES used as ’inspired by’ leads. These examples teach the model the expected input-output format and the style of molecules we are looking for. When we then provide the model with our specific target pocket description, it generates candidate molecules as SMILES strings, one character at a time. The transformer self-attention mechanism allows it to consider the entire context simultaneously, including both the prompt examples and the SMILES being generated, thereby helping maintain chemical coherence (e.g., ensuring that rings close properly, parentheses balance, and bond symbols are valid).

### 2.3 Experimental Data and Small Molecule Selection

In this study, we utilize a chemically diverse set of small-molecule inhibitors targeting ACVR1 (Activin A receptor type I). These include high-potency chemical probes and pre-clinical leads optimized for selectivity over other TGF-*β* family receptors (see Table 2).

**Table 2:**
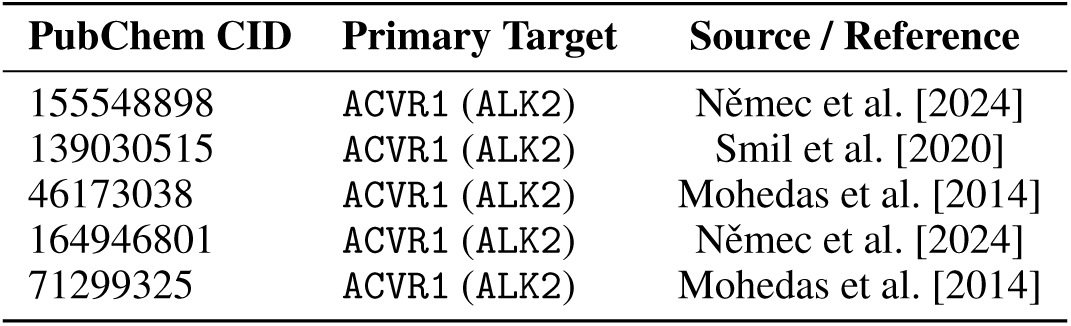
Selected small molecule inhibitors targeting ACVR1 (ALK2).

| PubChem CID | Primary Target | Source / Reference |
| --- | --- | --- |
| 155548898 | ACVR1 (ALK2) | Němec et al. [2024] |
| 139030515 | ACVR1 (ALK2) | Smil et al. [2020] |
| 46173038 | ACVR1 (ALK2) | Mohedas et al. [2014] |
| 164946801 | ACVR1 (ALK2) | Němec et al. [2024] |
| 71299325 | ACVR1 (ALK2) | Mohedas et al. [2014] |

Moreover, to target the B-cell receptor (BCR) signaling pathway associated with CD19, we select a panel of clinically approved and investigative inhibitors. This set includes first and second-generation covalent BTK inhibitors, BCL-2 inhibitors, and PI3K*δ*-selective agents. These molecules represent the standard of care for disrupting survival signals in CD19-expressing B-cell lymphomas (see Table 3).

**Table 3:**
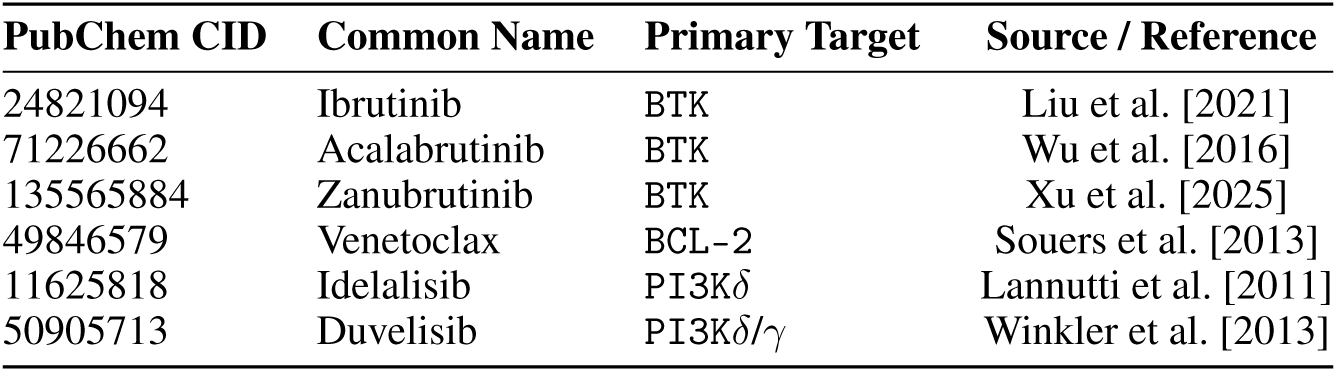
Panel of clinically approved and investigative inhibitors of BCR.

### 2.4 Biophysical Evaluation with Boltz-2

After the generation phase, for each candidate molecule created by the LLM, we perform a rigorous affinity evaluation to predict its binding orientation and strength. We use Boltz-2 to predict the bound complex structures and affinity metrics [Passaro et al., 2025]. Boltz-2 treats the protein-ligand system as a graph of atoms and employs a diffusion module to arrange them into a low-energy state. This process is guided by a neural network that has learned the physics of biomolecular interactions from millions of known complexes [Jumper et al., 2021, Ho et al., 2020]. This approach acts as a high-speed computational surrogate for molecular dynamics, outputting plausible bound poses and confidence measures in seconds. Boltz-2 was run in single-sequence mode without a multiple sequence alignment, and without physics-based potentials.

We provide Boltz-2 with our protein structure and the generated ligand SMILES strings. To improve accuracy, we constrain the search by specifying the binding pocket residues. Boltz-2 then predicts a protein-ligand complex structure and an affinity probability in the range [0, 1]. The model also outputs confidence scores similar to AlphaFold, such as per-residue confidence (pLDDT) and interface confidence (ipTM).

We filter the results by rejecting any candidate that failed to produce a clear pocket-bound pose or yielded low confidence scores. For successful poses, we record two independent Boltz-2 outputs: the affinity probability and the predicted potency (p*IC*_50_); *IC*_50_ is derived from p*IC*_50_ alone (*IC*_50_ = 10*^−^*^p^*^IC^*^50^ M). We prioritize complexes with an ipTM *>* 0.95, as this indicate a reliable binding mode. Because the Boltz-2 diffusion approach implicitly accounts for protein flexibility, it captures induced-fit effects that traditional docking might miss, offering accuracy comparable to intensive physics-based simulations [Raschka et al., 2018, Wang et al., 2019].

Finally, Boltz-2 serves dual purposes by filtering out molecules with poor or weak binding potential and ranking the remaining candidates by predicted affinity and structural quality. For each target, we typically advance only the top 5–10 candidates showing the highest predicted affinity and no structural anomalies.

### 2.5 Iterative Feedback-driven Optimization

Candidate molecules undergo further refinement through a feedback-guided iterative optimization loop informed by Boltz-2 outputs and cheminformatic scoring. The LLM acts as an agent that progressively improves its proposals as it is fed structured feedback on previous generations. A running registry accumulates every molecule that clears the scoring gates across all iterations, together with its Boltz-2 affinity, structural confidence, synthetic accessibility (SAS), and novelty metrics; molecules already scored in an earlier iteration are served from a cache.

At each new iteration the prompt is rebuilt from three pools: Positive leads—a fixed pair of anchor seed molecules plus the highest-scoring, structurally distinct candidates found so far. A candidate is only admitted to this pool if its Tanimoto similarity to every lead already present is below 0.9, which keeps the feedback pool from collapsing onto one scaffold. Negative examples are split into two contrastive failure modes so the model can tell why a molecule failed: molecules that bound strongly but were too hard to synthesize, and molecules that were easy to synthesize but bound too weakly. Already-proposed molecules from the current run, so the model avoids emitting exact duplicates. This yields prompts such as: "Generate analogs of [leads]. AVOID repeating the failure patterns of strong binders that were too hard to synthesize, like [X]; and easily synthesizable molecules that bound too weakly, like [Y]. Already proposed, do not repeat: [Z]."

Molecular performance is quantified using a simple reward function that combines the Boltz-2 affinity score as the primary component, with a novelty term that encourages structural diversity and prevents the model from repeatedly generating similar molecules. The total reward *R*(*m*) *∈* [0, 1] for a generated molecule *m* is defined as:

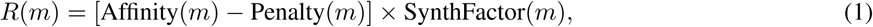

Molecules with an SAScore above *SAS_max_* = 6 are excluded from the candidate pool prior to scoring; among the remaining candidates, SynthFactor(*m*) scales the reward toward zero as *SAS*(*m*) approaches this exclusion threshold, so among molecules with similar affinity, easier-to-synthesize candidates are favored:

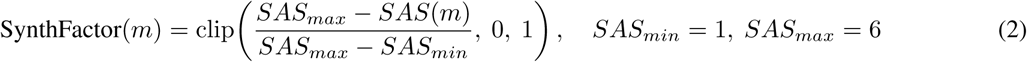

where the affinity term is derived from the Boltz-2 predicted binding probability. We apply a threshold to prioritize high-confidence candidates:

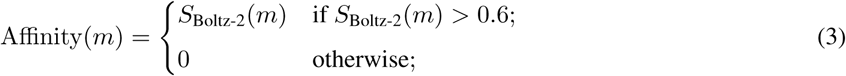

where *S*_Boltz-2_(*m*) *∈* [0, 1] denotes the Boltz-2 binding probability score for molecule *m*.

To prevent mode collapses and ensure exploration of novel chemical space, we apply a penalty to redundant structural analogs. If the maximum Tanimoto similarity to any entry in the global registry *G* exceeds 0.9, the reward is reduced by 50%:

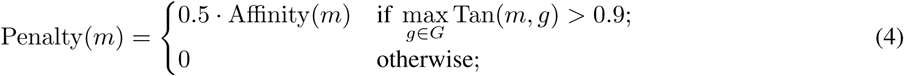

where Tan(*m, g*) *∈* [0, 1] denotes the Tanimoto similarity between molecule *m* and registry entry *g*, calculated using Morgan fingerprints (using 2048 bits and radius = 2). Over successive iterations, this approach drives convergence toward consistently high-score proposals. To prevent premature loss of chemical diversity, we implement a similarity filter. When a newly generated molecule shows high structural similarity to an existing top candidate (Tanimoto similarity *>* 0.9), we reduce its reward. This ensures that each iteration continues to explore new chemical structures rather than simply repeating successful ones.

### 2.6 Cheminformatic Filtering and Validation

Before selecting any compound as a potential lead, we subject all candidates to post-generation cheminformatics validation to ensure that our AI-generated molecules met standard criteria for drug-likeness and synthetic feasibility. We calculate the Quantitative Estimate of Drug-likeness [Bickerton et al., 2012] (QED) for each molecule using RDKit [RDKit, 2024], an open-source cheminformatics Python library. QED represents a metric ranging from 0 to 1, where higher values indicate that molecular properties such as molecular weight, LogP (octanol-water partition coefficient), and topological polar surface area fall within ranges typical of approved oral drugs. This metric goes beyond simple yes/no rules by capturing how different properties contribute to drug similarity.

We subsequently compute the Synthetic Accessibility Score [Ertl and Schuffenhauer, 2009] (SAScore) for each molecule, an estimator that combines fragment rarity and structural complexity to predict synthetic difficulty. Scores range approximately from 1, representing very straightforward synthesis such as benzene, to 10, indicating considerable synthetic challenges. We establish a threshold by which molecules with SAScores greater than 6 warrant careful scrutiny or exclusion [Ertl and Schuffenhauer, 2009], as values above this threshold typically suggest significant synthetic difficulties involving multiple steps or exotic reagents. Using substructure searches in RDKit, we filter out any molecules containing Pan-Assay Interference Compounds (PAINS) motifs [Dahlin et al., 2015, Lagorce et al., 2015] as defined by the known alerts compiled by Baell and Holloway [Baell and Holloway, 2010] or BRENK substructure alerts, as implemented in RDKit. These problematic scaffolds include catechols, curcumin-like enones, aggregators, and other structures frequently associated with false-positive results. Such compounds often demonstrate activity in screening assays through artifact mechanisms rather than genuine targets [McGovern et al., 2002]. All final candidates are free of PAINS alerts and of obviously reactive functional groups such as alkyl halides or Michael acceptors. This filtering step helps avoid molecules that look good computationally but fail in the lab due to poor properties or false positive signals in assays [Chen et al., 2018, Vamathevan et al., 2019].

### 2.7 Patent and Novelty Assessment

To ensure that our de-novo designed molecules were truly novel and unencumbered by existing patents, we perform a novelty screen. For each top candidate SMILES, we query the PubChem [Kim et al., 2025] database via its API to see if an identical structure exists (checking for an assigned PubChem CID). If a match was found, we examine the associated records to identify if the compound was known as a drug, an experimental tool compound, or mentioned in patent literature. All of our final leads returned no match in PubChem, indicating they are likely novel chemical entities (NCEs). This step is important for gauging freedom-to-operate and also simply to validate that the LLM has not memorized a known drug. These results increase our confidence that LLMsFold is indeed exploring new regions of chemical space rather than rediscovering known molecules.

### 2.8 Computational Implementation

The computational architecture has been built as an asynchronous Python framework querying several API endpoints. One endpoint is represented by the Groq^4^ inference engine, which provides quick access to the Llama-3-70B model for high-frequency SMILES generation, which initiates the process. After generation, the pipeline switches to the endpoint represented by the Boltz-2 foundation model via the NVIDIA Cloud Functions (NVCF)^5^ interface. The asynchronous nature of the architecture, made by using Python asyncio and httpx libraries, renders the pipeline very efficient, allowing PubChem novelty checks to run concurrently across candidates; Boltz-2 evaluations are currently submitted sequentially. For all other chemical calculations, the pipeline uses RDKit to compute molecular properties (LogP, QED, and SAS) that guide the iterative optimization process, as well as molecular fingerprints for similarity comparisons.

### 2.9 Proteins and Ligands Visualization

Three-dimensional visualization of target proteins (ACVR1 and CD19) and small-molecule ligands has been performed using UCSF ChimeraX^6^ [Pettersen et al., 2021]. UCSF ChimeraX is a molecular visualization and analysis tool developed by the Resource for Biocomputing, Visualization, and Informatics (RBVI), University of California, San Francisco. Small molecules generated in this study are not shown or described in this article, as they are intended for patent protection pending experimental validation.

## 3 Results

Applying LLMsFold to the ACVR1 and CD19 targets yields a set of novel small-molecule inhibitor candidates that successfully passed all stages of our pipeline. In total, we generate 50 unique molecules for each target, from which a handful of top candidates (n=15) emerged after iterative refinement. Table 4 and Table 5 summarize key properties of the leading compounds for each target pocket. These molecules are drug-like (molecular weight *∼* 400*/*500, QED, SAS, predicted *IC*_5_0, etc.) and novel (no PubChem hits for patented drugs). We describe the results for each target in detail below.

**Table 4:** Computational binding predictions and drug-likeness metrics for ACVR1 lead candidates. Note that *pIC*_50_ and *IC*_50_ values are predicted by the Boltz-2 model.

| Molecule | MolWt | QED | SAS | $pIC_{50}$ | $IC_{50}$ | Aff. Prob |
| --- | --- | --- | --- | --- | --- | --- |
| Molecule 1 | 448.57 | 0.590 | 2.64 | 10.72 | 0.019 nM | 0.953 |
| Molecule 2 | 419.53 | 0.637 | 2.81 | 10.51 | 0.031 nM | 0.949 |

**Table 5:** Computational binding predictions and drug-likeness metrics for CD19 lead candidates. Note that *pIC*_50_ and *IC*_50_ values are predicted by the Boltz-2 model.

| Pocket | MolWt | QED | SAS | $pIC_{50}$ | $IC_{50}$ | Aff. Prob |
| --- | --- | --- | --- | --- | --- | --- |
| Pocket 1 | 433.27 | 0.519 | 2.57 | 7.73 | 18.6 nM | 0.540 |
| Pocket 2 | 460.25 | 0.563 | 2.86 | 5.43 | 3.75 μM | 0.367 |
| Pocket 3 | 331.15 | 0.597 | 2.34 | 6.51 | 311.2 nM | 0.160 |

### 3.1 ACVR1 (Activin A Receptor Type 1)

ACVR1 (Activin A Receptor Type 1) is a type I BMP receptor belonging to the TGF-*β* superfamily (PDB: 6ACR). The R206H mutation causes fibrodysplasia ossificans progressiva (FOP), a rare disorder where soft tissues progressively turn into bone. This mutation makes the receptor constitutively active, triggering aberrant osteogenic signaling even without ligand binding. ACVR1 is a serine/threonine kinase, and we targeted the ATP-binding pocket located between the N- and C-lobes of the kinase domain, aiming to block its hyperactive signaling [Hino et al., 2015]. LLMsFold generated several polycyclic heterocycle scaffolds reminiscent of known kinase inhibitors. For ACVR1, Boltz-2 identified 15 molecules from approximately 50 LLM-generated proposals with strong binding probability (affinity *>* 0.6) and high-confidence poses (ipTM *≈* 0.95, ligand pLDDT *>* 0.9), which were advanced to final refinement. The top two candidates are referred to as ACVR1 Molecule 1 and ACVR1 Molecule 2.

Figure 2 illustrates the predicted binding pose of Molecule 1 within the kinase domain of ACVR1 using a surface representation. The ligand is predicted to occupy the ATP-binding pocket, forming a potential hydrogen bond with residues lining the catalytic cleft. The R206H mutation, located in the regulatory GS domain adjacent to the kinase region, is known to destabilize the inactive conformation of ACVR1, leading to increased pocket accessibility and altered kinase regulation [Hino et al., 2015]. In this configuration, the active site is largely enclosed by the ligand. Boltz-2 predicts a very high binding confidence for ACVR1 Molecule 1: interface TM-score ipTM = 0.983 (indicating an almost exact alignment of key interface contacts) and an average ligand pLDDT of 0.958, meaning the ligand position is very well-defined [Wohlwend et al., 2024]. The affinity probability was 0.953, corresponding to a predicted *pIC*_50_ *≈* 10.72 (*IC*_50_ *∼* 1.9 *×* 10*^−^*^11^ M, or 0.019 nM). This is an encouraging potency for an initial design. Four distinct binding poses were output by Boltz-2.

**Figure 2:**
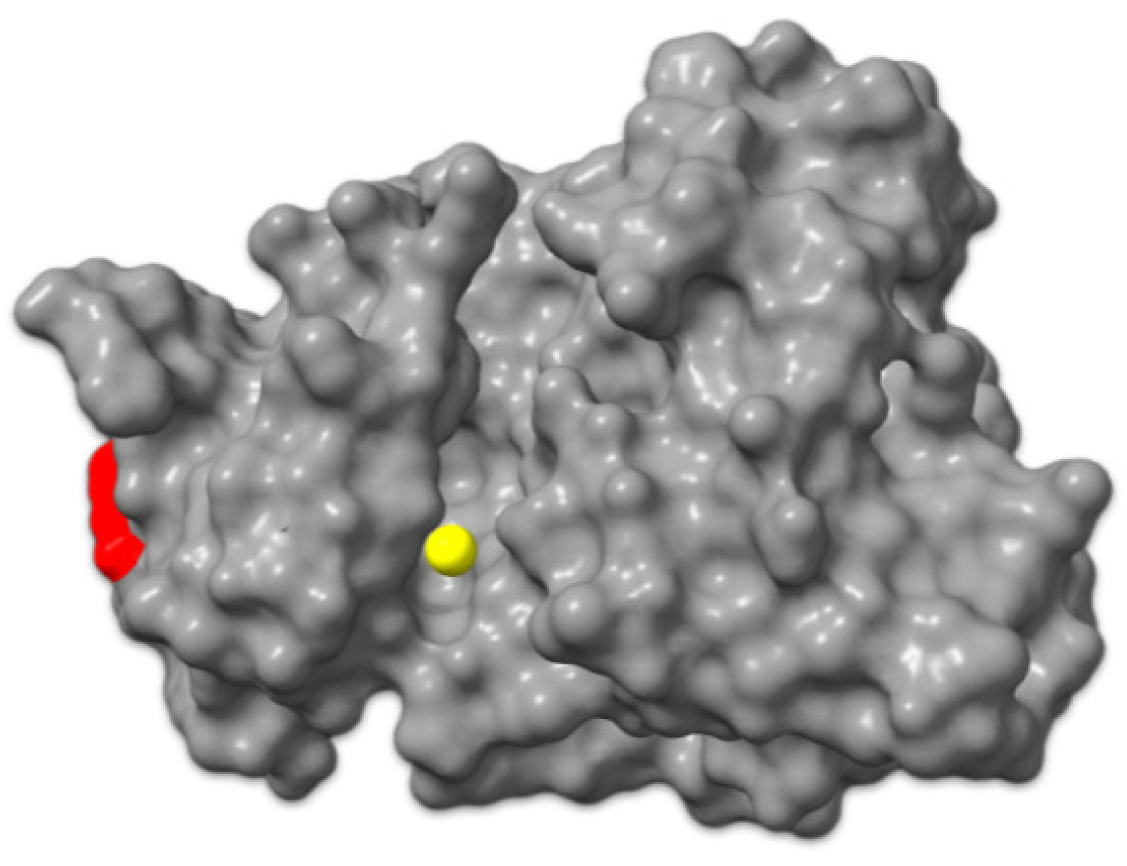
Surface view of ACVR1 pocket and its ligand. Molecular surface representation of the ACVR1 kinase domain (gray) showing the ATP-binding cleft occupied by the LLMsFold-generated ligand (yellow). The R206 residue (red), site of the R206H substitution causing constitutive kinase activation in fibrodysplasia ossificans progressiva (FOP), is located adjacent to the binding pocket.

We also identified ACVR1 Molecule 2, an analog of Molecule 1. It shows similarly high confidence (ipTM 0.983) with similar affinity probability (0.949, *pIC*_50_ *≈* 10.51, *∼* 0.031 nM). Nonetheless, Molecule 2 provides an alternative scaffold, and its bromine could be useful for future medicinal chemistry (as a handle for substituent variation). ACVR1 leads have QED scores ranging from 0.55 to 0.64, indicating moderate to good drug-likeness and suggesting favorable physicochemical properties consistent with oral bioavailability. They satisfy all four Lipinski criteria, which are enforced as a hard filter in the pipeline. Their Synthetic Accessibility (SA) scores are 2.64 and 2.81, respectively, meaning they are considered easy-to-moderate to synthesize; indeed, they resemble multi-ring scaffolds accessible via known coupling reactions. We have verified that no PAINS motifs are present. Thus, ACVR1 candidates are potent, well-fitting, and chemically reasonable starting points for inhibitor development. Given the role of ACVR1 in Fibrodysplasia Ossificans Progressiva (FOP) and certain cancers [Shore et al., 2006], these compounds (if experimentally confirmed) could represent valuable leads.

### 3.2 CD19

CD19 is a type I transmembrane glycoprotein (PDB: 7URV) expressed throughout B-cell development, from early pro-B cells to mature B lymphocytes, but lost upon plasma cell differentiation [Wang et al., 2012]. It functions as a co-receptor within the B-cell receptor signaling complex, associating with CD21, CD81, and Leu-13 to lower the activation threshold for antigen-mediated signaling [He et al., 2023]. Given its consistent and high expression on B-cell malignancies, CD19 has become a key therapeutic target: it is the antigen recognized by blinatumomab (a bispecific T-cell engager), tafasitamab (an Fc-enhanced monoclonal antibody), and several approved CAR-T therapies including tisagenlecleucel and axicabtagene ciloleucel [Li et al., 2017]. These approaches rely on extracellular recognition by large biologic therapies. These grooves could potentially accommodate small molecules designed to disrupt protein–protein interactions within the CD19 signaling complex. For CD19, we identified a total of 7 molecules across three pockets, with binding affinities ranging from 0.1 to 0.6.

Pocket 1 is located at the membrane-proximal interface of the CD19 ectodomain. Since CD19 does not contain a deep catalytic cavity, we first anchored our analysis to a biologically validated interaction site. Using the available cryo-EM structure of CD19 in complex with the FMC63 antibody fragment (PDB: 7URV) [He et al., 2023], whose scFv domain is used in approved CAR-T therapies, we identified two shallow pockets (Pocket 1 and Pocket 2) at the membrane-proximal domain interface. This structure defines a clinically relevant epitope on the protein surface (Fig. 3). We then evaluated whether LLMsFold molecules bind near this region. Boltz-2 predicted a stable pose for the top candidate within Pocket 1 (Fig. 4). The ligand binds adjacent to and partially overlaps the antibody epitope, suggesting possible interference with protein–protein recognition. As visible by comparing Fig. 3 and Fig. 4, the predicted molecule partially occupies the same functional surface region. The predicted binding probability was 0.54 and the estimated *pIC*_50_ was 7.73 (*IC*_50_ *≈* 18.6 nM). Considering the shallow surface, this indicates a favorable binding for a protein–protein interaction site. Overall, the predicted pose supports the biological relevance of CD19 Pocket 1 molecule and suggests that the generated molecules target a functional interaction interface rather than a random surface region.

**Figure 3:**
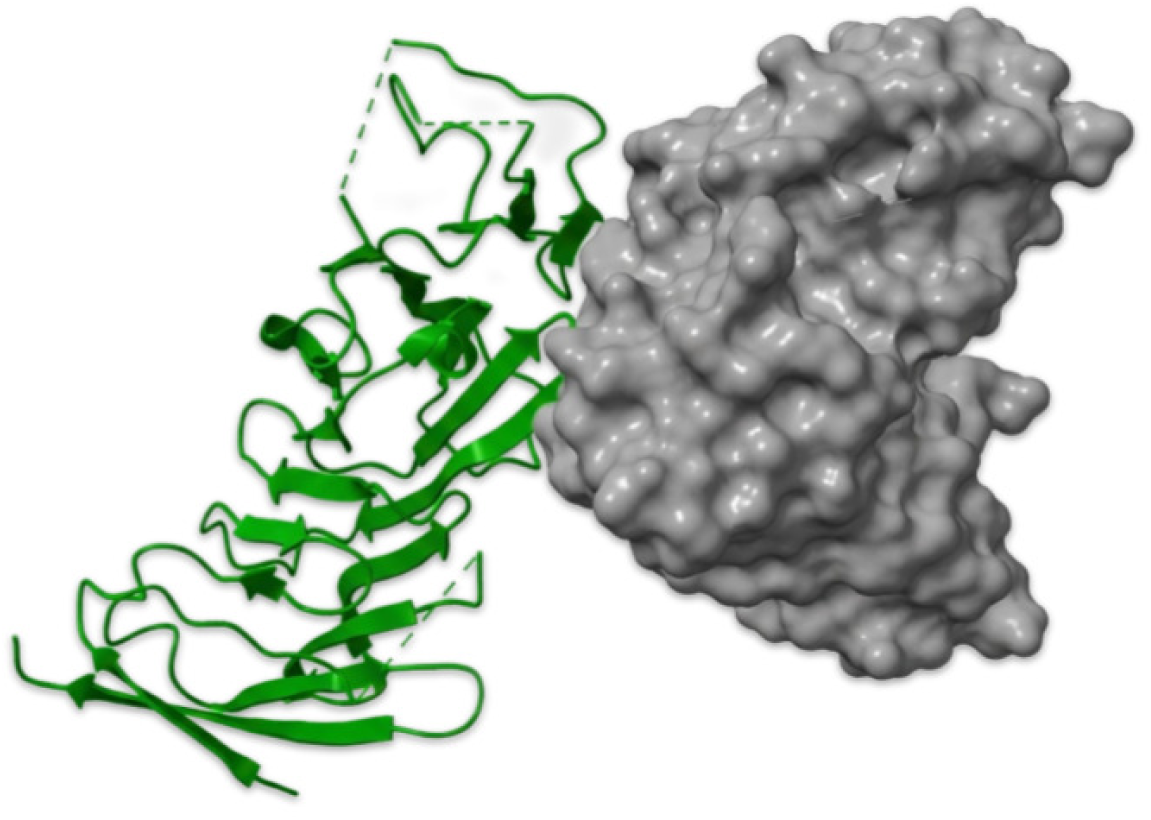
Experimental CD19–FMC63 complex. CD19 is shown as a molecular surface (gray) and the FMC63 scFv in green. This interaction site was used to identify Pocket 1.

**Figure 4:**
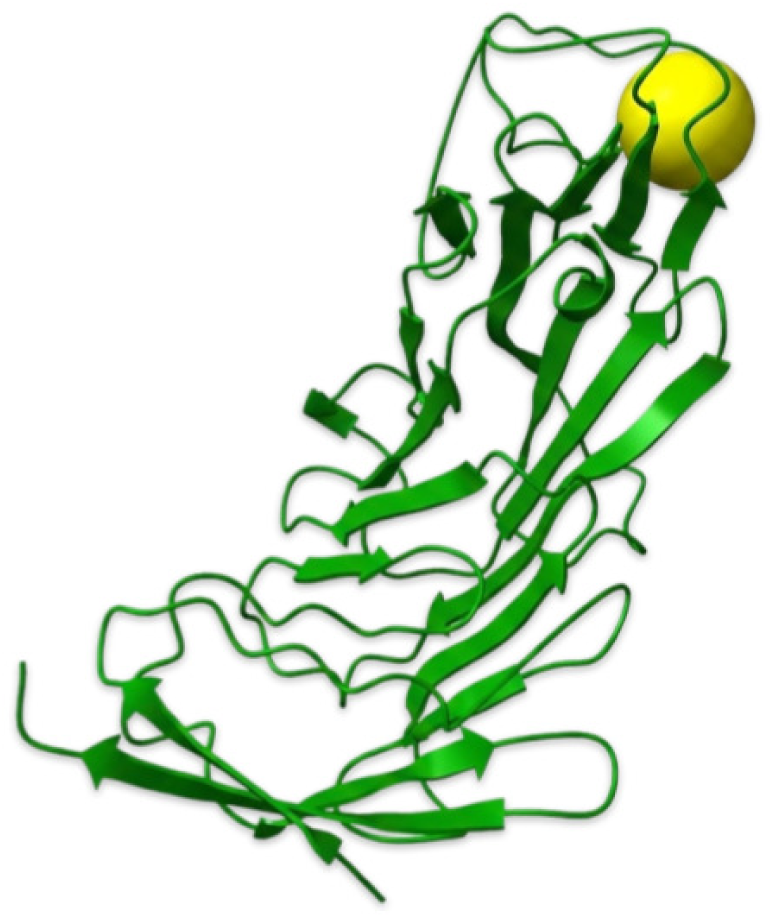
Predicted binding pose of the lead small-molecule candidate in CD19 Pocket 1. CD19 is shown in green as a cartoon representation. The molecule binds close to the FMC63 epitope shown in Figure 3. The ligand structure is not disclosed for intellectual property protection.

Pocket 2 is a slightly deeper cavity located at a domain interface on CD19, on the opposite face of the protein compared to Pocket 1 (Figure 5). For this site, LLMsFold tended to generate larger and more polar molecules with a somewhat peptide-like character, which reflects the different chemical environment of this pocket. The best candidate for this pocket has been predicted by Boltz-2 with high interface confidence (ipTM = 0.931) and adopted a pose that bridges the two sides of the pocket. However, the predicted binding affinity was weaker than for Pocket 1, with an affinity probability of 0.37 corresponding to a *pIC*_50_ of approximately 5.43, or an *IC*_50_ of about 3.7 *µ*M. Despite the lower predicted affinity, the docking score (*−*6.695) was actually better than those obtained for Pocket 1, suggesting a favorable geometric fit. The molecule also showed reasonable drug-like properties, with a molecular weight of approximately 460 Da, a QED score of 0.56, and a synthetic accessibility score of 2.86 (Table 5). While micromolar potency is modest, it represents a viable starting point for further optimization, particularly for a novel protein-protein interaction inhibitor targeting a site distinct from the FMC63 epitope.

**Figure 5:**
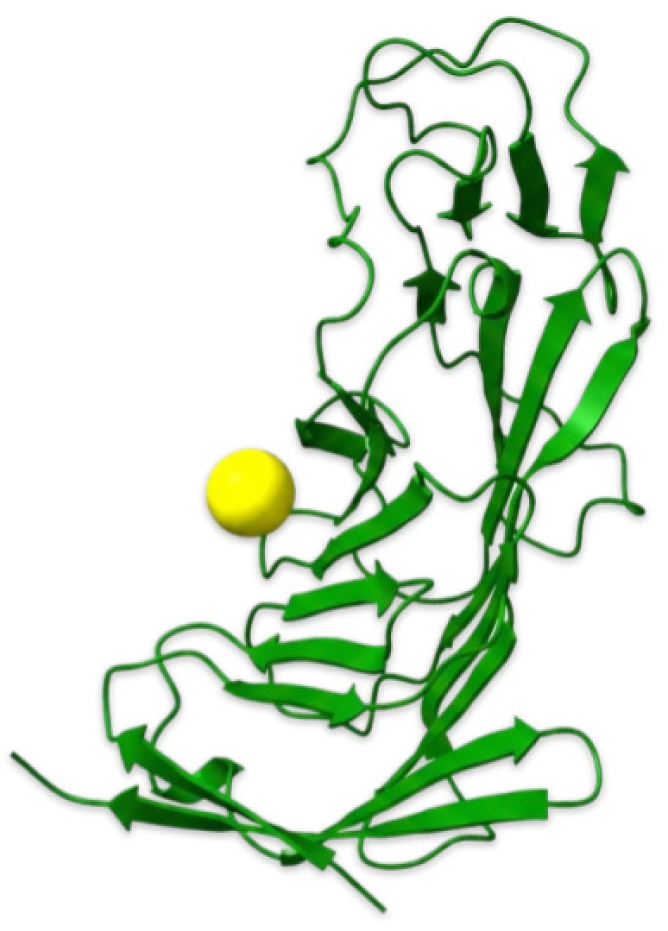
Predicted binding pose for Pocket 2 on CD19. The protein is shown as a green cartoon, and the ligand position is represented as a yellow sphere. Unlike Pocket 1, this binding site is located on the opposite face of the protein, away from the FMC63 epitope.

We also found a Pocket 3 on CD19 (a shallow surface groove). One interesting molecule had a predicted *pIC*_50_ *∼* 6.5 (*≈* 300 nM). While Boltz-2 gave it moderate confidence (ipTM *∼* 0.90), its affinity probability was only 0.16; thus, we consider it a lower-priority hit. It also had above-average drug-likeness (QED *∼* 0.597).

### 3.3 Validation and Comparisons

All top molecules from ACVR1 and CD19 were checked for novelty as described: none were in PubChem or had known activity, underscoring the capability of the LLM in generating de novo structures rather than retrieving known ones. From a medical and chemical perspective, the ACVR1 Molecule 1 is perhaps the most compelling because it is sufficiently distinct from any single known inhibitor to be considered novel.

Finally, we compared our Boltz-2-predicted binding modes with classical docking to provide a reality check. We ran AutoDock Vina [Eberhardt et al., 2021, Trott and Olson, 2010] on ACVR1 with Molecule 1, and it produced a pose very similar to the Boltz-2 top pose (RMSD *<* 1.5 Å) with a docking score of *−*6.2 (Vina units). This corroborates that Molecule 1 is indeed likely to bind in the intended pocket. For CD19, classical docking is less reliable due to the shallow nature of the pockets, and indeed Vina struggled (producing very diffuse poses). This is where the strength of Boltz-2 lies because it can handle cases beyond the realm of rigid docking.

A benchmarking comparison against State-of-the-Art approaches — Pocket2Mol Peng et al. [2022], REINVENT 4 Blaschke et al. [2020], Loeffler et al. [2024], S3-GFN Kim et al. [2026], and SynFlowNet Cretu et al. [2025] — showing that LLMsFold achieves the strongest predicted affinities, is reported in Appendix A.

## 4 Discussion

The central design principle of LLMsFold is the integration of structural awareness into every stage of molecular generation. Unlike conventional virtual screening, where large pre-enumerated libraries are filtered post hoc, LLMsFold conditions generation on the residue composition of the target binding site. The pipeline begins with an unbiased, geometry-based identification of druggable pockets and feeds this spatial information directly into the language model prompt, constraining the generative process to a biologically meaningful region of the protein surface. Boltz-2 then serves a dual function: it evaluates each candidate by predicting the three-dimensional bound complex and associated confidence metrics, and it feeds this evaluation back into the generative loop, where successful molecules are reintroduced as prompt examples for subsequent rounds. This creates an optimization trajectory in which the language model progressively adapts its proposals across iterations, converging toward scaffolds complementary to the pocket geometry. The process thus resembles a guided optimization cycle rather than a single-pass sampling procedure, distinguishing LLMsFold from classical docking-then-filtering workflows.

A distinctive methodological choice is the exclusive reliance on in-context learning rather than fine-tuning [Dong et al., 2024]. Most existing generative frameworks for drug design, including REINVENT [Olivecrona et al., 2017] and DrugGPT [Li et al., 2023], require task-specific training on large chemical datasets, demanding substantial computational resources and curated corpora. LLMsFold instead exploits the chemical knowledge already embedded in the pre-trained weights of Llama-3-70B, acquired during large-scale pretraining on diverse text that includes scientific literature and chemical notation [Touvron et al., 2023, Brown et al., 2020]. This strategy eliminates the need for dedicated GPU clusters and weeks of fine-tuning, and it enables rapid target switching: adapting the pipeline to a new protein requires only reformulation of the prompt with appropriate pocket descriptors and reference inhibitors. Scaling laws for large language models further suggest that in-context learning quality improves with model size [Kaplan et al., 2020, Wei et al., 2022], implying that future, larger models may enhance generation quality without architectural changes to the pipeline. At the same time, in-context learning carries inherent constraints. Increasing the number of reference molecules in the context can lead to deteriorated performance, as larger contexts often introduce additional noise and complexity. Moreover, the absence of gradient-based optimization limits the model’s ability to deeply internalize structure and activity relationships for a specific target.

The CD19 results illustrate a particularly challenging application. Unlike ACVR1, which presents a well-defined ATP-binding cleft typical of kinases, CD19 lacks deep catalytic cavities. Its druggable surface consists of shallow grooves at domain interfaces, a topology that has historically resisted small-molecule intervention. Protein–protein interaction (PPI) surfaces are among the most difficult targets in drug discovery: they are flat, solvent-exposed, and extend over large areas, making it hard for small molecules to achieve sufficient binding affinity [Wells and McClendon, 2007, Arkin et al., 2014]. In this context, the ability of LLMsFold to propose molecules with sub-micromolar predicted affinity for CD19 Pocket 1 (*pIC*_50_ *≈* 7.73) is noteworthy. The predicted pose overlaps with the FMC63 antibody epitope partially, suggesting possible interference with a clinically validated interaction surface, consistent with the emerging strategy of targeting antibody-binding epitopes with small molecules as an alternative to biologics [June and Sadelain, 2018]. The performance gradient across the three CD19 pockets provides additional insight: Pocket 1, overlapping the FMC63 epitope, yielded the highest predicted affinity, consistent with the notion that functionally constrained surfaces present more defined pharmacological features. Pocket 2 produced candidates with micromolar affinity but favorable docking scores, suggesting geometric complementarity amenable to focused optimization. Pocket 3 yielded a lower-confidence hit. This differential performance may guide future efforts toward the most tractable binding sites and illustrates how LLMsFold can serve as a tool for druggability assessment of novel protein surfaces.

A practical finding is that the entire pipeline executes in minutes and requires no local GPU, since molecular generation and co-folding are executed on remote API endpoints; the client can be an ordinary laptop. This accessibility has direct implications for rare and neglected diseases, where research funding and computational infrastructure are often limited. Fibrodysplasia ossificans progressiva, caused by the ACVR1 R206H mutation, affects approximately one in two million individuals worldwide [Shore et al., 2006]. The extreme rarity of the condition means that pharmaceutical investment in traditional drug discovery campaigns is constrained by unfavorable market incentives [Tambuyzer et al., 2020]. Tools that reduce the cost and time of early-stage lead identification could therefore have a disproportionate impact on orphan drug development, lowering the barrier to entry for academic groups, patient foundations, and small biotech companies that lack access to high-performance computing clusters.

A few limitations should be considered. All binding affinities and structural predictions reported here are computational estimates. Boltz-2, while demonstrating accuracy approaching that of AlphaFold3 on established benchmarks [Passaro et al., 2025, Abramson et al., 2024], remains a predictive model whose outputs require experimental confirmation through biophysical assays such as surface plasmon resonance or isothermal titration calorimetry. The pipeline does not currently incorporate predictions of absorption, distribution, metabolism, excretion, and toxicity properties; a molecule with excellent predicted binding may nonetheless fail in vivo due to poor bioavailability or off-target toxicity. While QED and SA scores provide a first approximation of drug-likeness and synthetic feasibility, they do not substitute for dedicated pharmacokinetic modeling. Language models are also susceptible to chemical hallucinations, generating SMILES strings that parse correctly but encode chemically implausible or unstable structures. Although multi-stage filtering mitigates this risk, it cannot eliminate it entirely. Furthermore, the quality of generated molecules depends on the selection of reference compounds provided in the prompt; a poorly chosen set of examples may bias the model toward a narrow region of chemical space. Finally, the Boltz-2 affinity predictions, while useful for ranking, should not be interpreted as quantitative *IC*_50_ values without calibration against experimental data for the specific targets examined.

Looking ahead, the most immediate priority is the experimental validation of the top candidates through in vitro binding assays and cellular functional studies. Integration of toxicity prediction models directly into the feedback loop would enable simultaneous optimization of binding affinity and pharmacokinetic properties. The incorporation of retrosynthetic planning tools could further constrain generation toward molecules with validated synthetic routes, bridging computational design and practical medicinal chemistry. From a modeling perspective, the pipeline is architecture-agnostic: as newer foundation models become available for both text generation and biomolecular structure prediction, they can be integrated without modifying the overall framework.

## 5 Conclusions

LLMsFold shows that large language models, when combined with accurate structural prediction, can serve as useful tools for early-stage drug design. The pipeline takes clinically relevant molecules as starting points, uses an LLM to generate structural variants, and relies on Boltz-2 to identify those with favorable predicted binding. This process completes in minutes and runs on accessible hardware. Our case studies on ACVR1 and CD19 produced novel candidates that satisfy standard drug-likeness criteria and show promising computational binding profiles. The compounds proposed by the LLM differ substantially from established inhibitors for these targets, which may offer advantages in terms of intellectual property and chemical optimization. These results are computational predictions, and experimental validation will determine their true value. We view LLMsFold not as a replacement for medicinal chemistry expertise, but as a tool that can accelerate the early ideation phase.

### Key Points

- LLMsFold is an AI pipeline for designing small-molecule drugs.
- It generates and optimizes candidates for protein targets using an LLM and Boltz-2.
- It shows promising results on ACVR1 and CD19.

## Acknowledgments

The manuscript was edited with assistance from Grammarly and large language models (LLMs) to improve clarity and style.

## Author Contributions

Wageesha Widuranga: Data curation, Investigation, Methodology, Software, Validation, Writing – original draft. Davide Rigoni: Supervision, Visualization, Writing – original draft, Writing – review & editing. Fabio Bove: Investigation, Methodology. Dario Righelli: Investigation, Methodology. Salvatore Romano: Data curation, Investigation, Methodology, Software, Visualization, Writing – original draft. Rosa Visone: Writing – review & editing. Marilena V. Iorio: Writing – review & editing. Marco Grassia: Data curation, Software. Giuseppe Mangioni: Data curation, Software. Pietro Lio: Project administration, Resources, Supervision. Cristian Taccioli: Conceptualization, Funding acquisition, Methodology, Project administration, Resources, Supervision, Writing – original draft, Writing – review & editing.

## Data Availability

The source code and input files for LLMsFold are available at https://github.com/tacclab/llmsfold. Generated molecules are withheld pending patent protection (see Competing Interests); run logs and result tables are available from the corresponding author on request.

## **A** Introduction

This section aims to compare LLMsFold with other State of the art approaches to computational drug design, Pocket2Mol Peng et al. [2022], REINVENT4^7^ Blaschke et al. [2020], Loeffler et al. [2024], S3-GFN Kim et al. [2026] and SynFlowNet Cretu et al. [2025]. All methods were evaluated under identical target and pocket conditions, specifically the ATP binding site of the ACVR1 kinase R206H mutant. While the biological context was held constant, the generated entities differ substantially in modality and underlying generative mechanism. Due to the substantial computational cost associated with training and adapting the other methods, the benchmarking experiments are conducted only on the ACVR1 target in this study.

Our approach, LLMsFold, is a structure aware, SMILES based LLM guided framework for small molecule generation that incorporates biophysical validation using Boltz-2 Passaro et al. [2025]. In contrast, Pocket2Mol Peng et al. [2022] addresses small molecule generation in the context of a 3D protein binding pocket by operating directly on molecular graphs and atomic coordinates, using an autoregressive, structure based assembly strategy rather than a SMILES based representation. REINVENT Blaschke et al. [2020], Loeffler et al. [2024], by comparison, is a sequence based de novo molecular design framework that typically represents molecules as SMILES strings and uses a recurrent neural network policy optimized through reinforcement learning to generate compounds with desired properties. Unlike graph or 3D based generative methods, REINVENT does not explicitly model spatial geometry during generation, but instead learns a distribution over molecular strings and steers generation toward user defined objectives via reward optimization. SynFlowNet Cretu et al. [2025], on the other hand, is based on GFlowNets Bengio et al. [2023], whose action space uses chemical reactions and purchasable reactants to sequentially build new molecules. By incorporating forward synthesis as an explicit constraint of the generative mechanism, it aims at bridging the gap between in silico molecular generation and real world synthesis capabilities. In this model, the protein pocket enters only indirectly through the reward function, which scores generated molecules by their predicted binding affinity, thereby guiding molecule generation without being explicitly modeled in the generative process itself. S3-GFN Kim et al. [2026]also builds on the GFlowNet framework but departs from SynFlowNet’s reaction template based construction rather than restricting the action space to predefined chemical reactions and building blocks, it generates molecules directly as SMILES strings, using a pretrained SMILES language model (GP MolFormer) as prior and GFlowNet post training (relative trajectory balance) to adapt it to a task specific reward. Importantly, unlike Pocket2Mol, REINVENT, S3-GFN and SynFlowNet, which require target specific training or fine tuning to be applied to the ACVR1 binding site, LLMsFold can be deployed directly without any additional model adaptation. This makes LLMsFold particularly advantageous in scenarios where rapid target deployment and reduced computational cost are essential.

## **B** Materials and Methods

### **B.1** Pipelines and candidate pool

All five pipelines were run independently against ACVR1, producing a combined pool of 1319 candidate molecules. Each candidate was scored for predicted binding affinity, physicochemical drug likeness, synthetic accessibility, and novelty against known or patented chemical matter. Candidates were retained if they satisfied molecular weight *≤* 500 Da, LogP *≤* 5, *≤* 5 hydrogen bond donors, *≤* 10 hydrogen bond acceptors, TPSA *≤* 140 Å^2^, *≤* 10 rotatable bonds, SAS *≤* 5, and QED *≥* 0.4, each pipeline’s own reported per property filtered pool does not itself enforce the affinity threshold, which is why pipelines such as Pocket2Mol and REINVENT show pool mean affinities well below 0.5, the affinity gate is applied only in the combined, cross pipeline cascade of Figure 6. For each pipeline we report the size of its property filtered pool (*N*), the mean and best predicted affinity probability, the mean affinity of its own top 5 candidates, the mean QED and synthetic accessibility score (SAS) of that top 5, the fraction of unique Bemis Murcko scaffolds in its filtered pool, and an intra pipeline diversity index defined as the mean pairwise Tanimoto similarity of its filtered pool over Morgan fingerprints, radius 2, 2048 bit, with lower values indicating a more internally diverse set.

**Figure 6:**
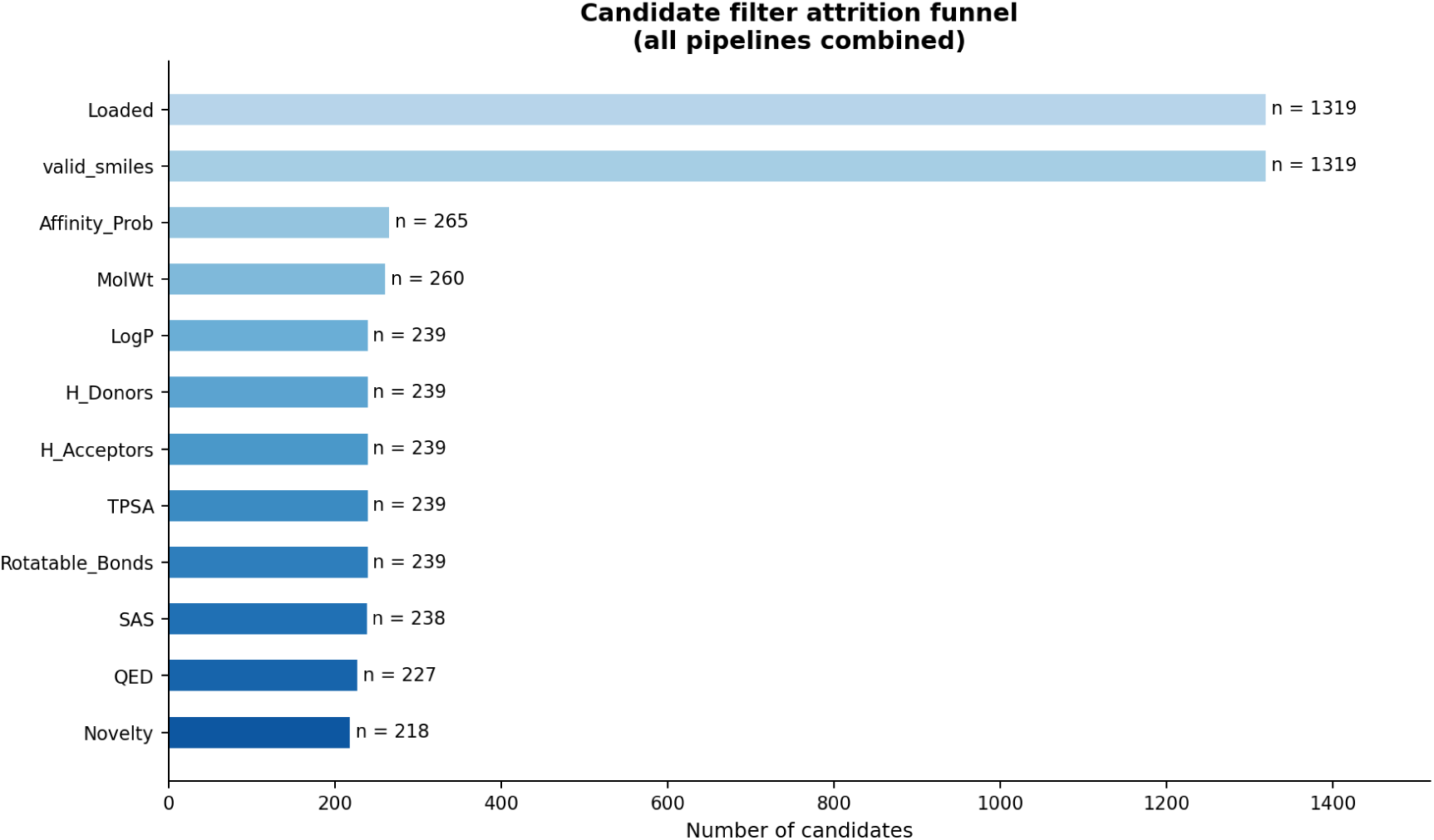
Filter attrition funnel, all five pipelines combined. All 1319 candidates parse as valid SMILES, the predicted affinity scoring stage (Affinity_Prob *≥* 0.5) reduces the pool to 265, molecular weight and LogP together remove a further 26 (260, then 239), four intervening property checks (hydrogen bond donors and acceptors, TPSA, rotatable bonds) cost nothing further, SAS removes 1, and QED and the final novelty check reduce 238 candidates to 218.

**Figure 7:**
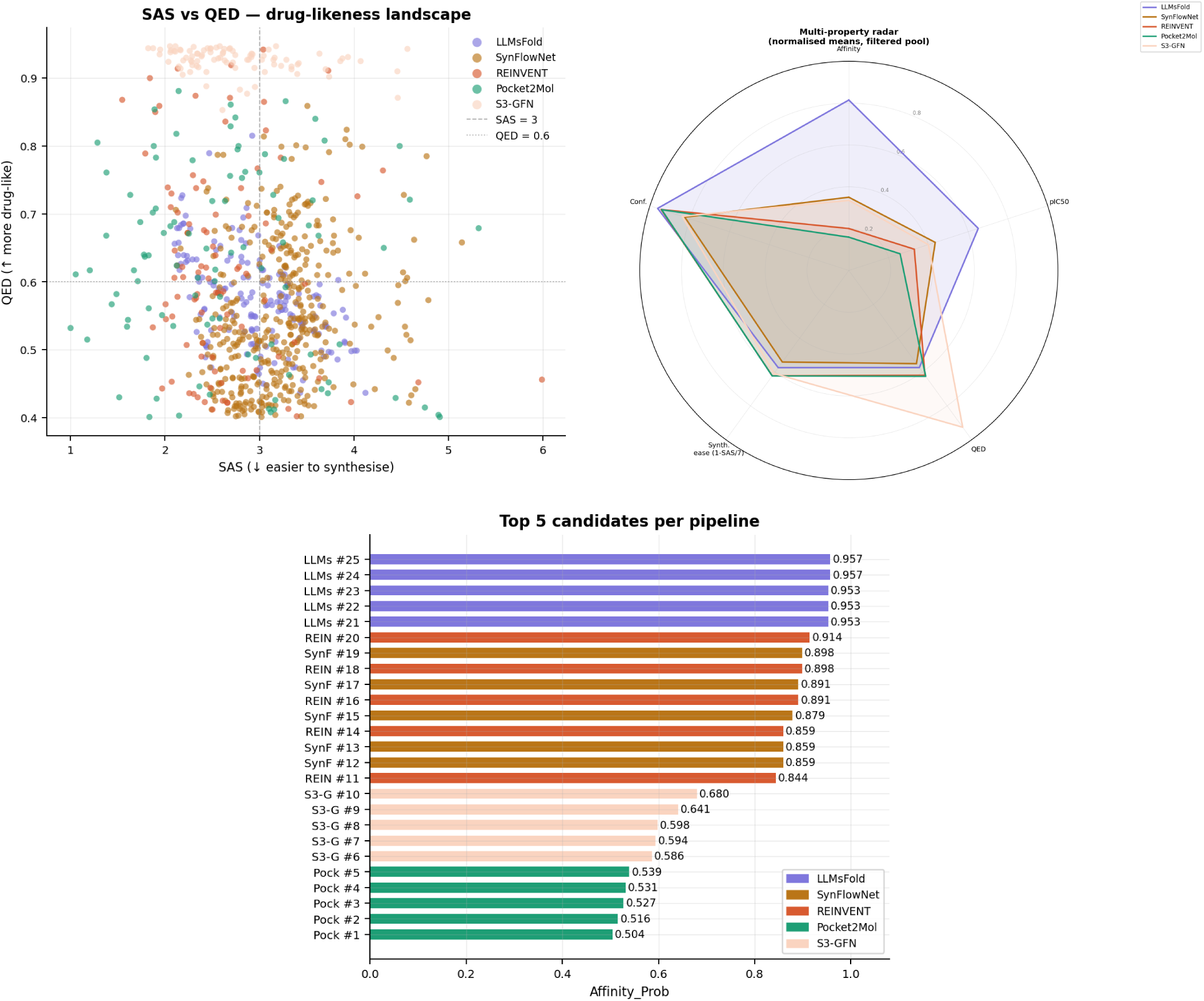
The top-5 affinity bar chart shows LLMsFold’s five best candidates clustered highest and tightest (0.945– 0.957), REINVENT’s and SynFlowNet’s best candidates interleave just below (0.84–0.91), S3-GFN’s top-5 sit in a middle band (0.59–0.68), and Pocket2Mol’s are lowest (0.50–0.54). This ranking of best candidates is a poor guide to typical output. The SAS versus QED landscape shows S3-GFN’s points (pale peach) clustering at the top of the QED axis across a wide range of SAS, distinct from LLMsFold, SynFlowNet and REINVENT’s overlapping central clustering and from Pocket2Mol’s low QED, high SAS spread.

**Figure 8:**
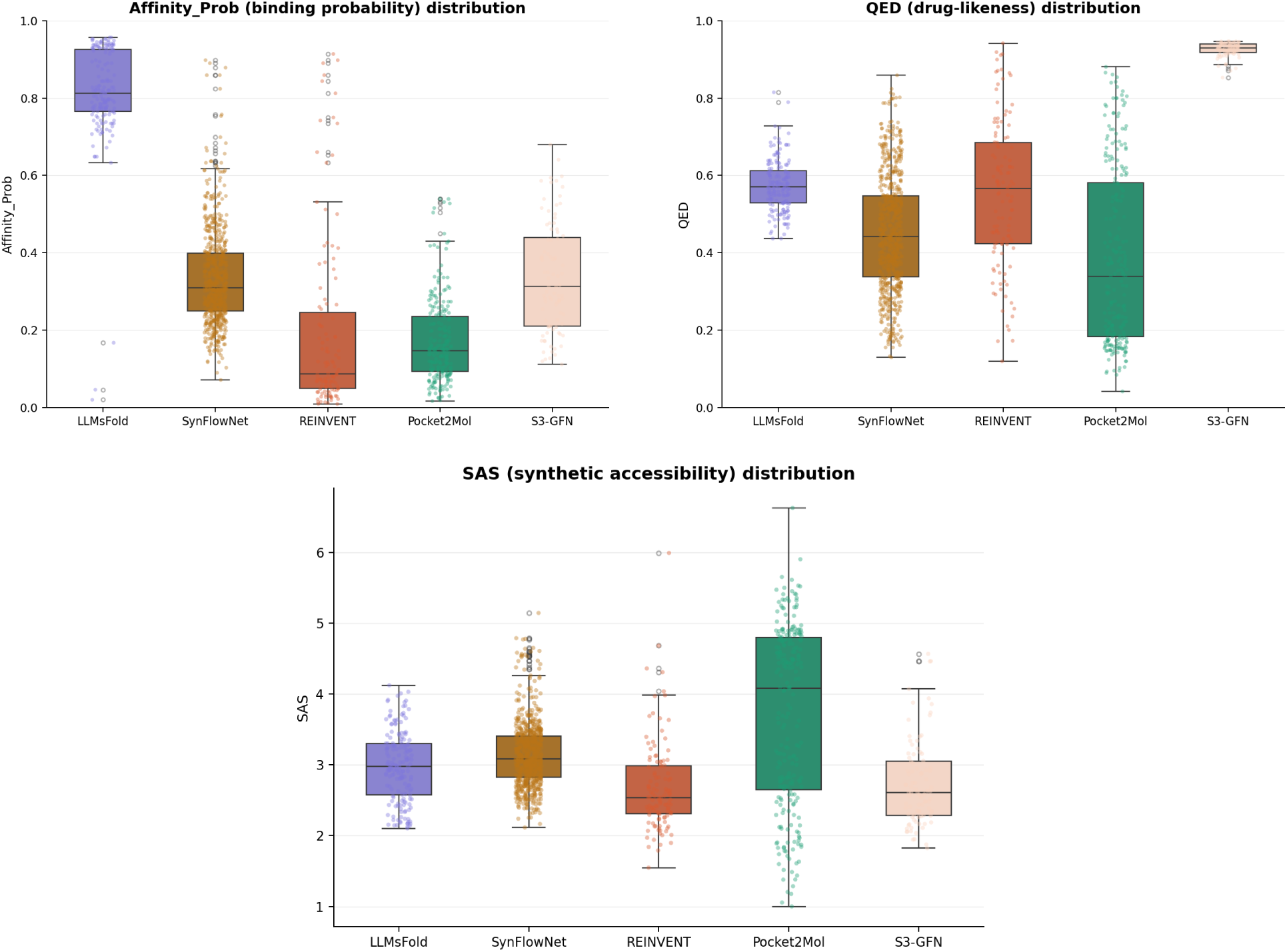
Per-molecule distributions over the full filtered pool (not just top 5), by pipeline. LLMsFold’s and Syn- FlowNet’s affinity distributions sit high and comparatively tight; REINVENT and Pocket2Mol show wide, right-skewed distributions with a long low-affinity tail; S3-GFN’s distribution sits in a middle range. QED (top right) shows S3-GFN’s entire interquartile range sitting above every other pipeline’s third quartile. SAS (bottom) highlights that Pocket2Mol’s median is the least favorable in the panel despite its top 5 having favorable scores (Table 6). whereas LLMsFold applies that full cascade inline during generation, so its count already reflects a post filter pool, and matches its *N* in Table 6 exactly.

### **B.2** Filtering cascade

All 1319 candidates parse as valid SMILES, so the first real attrition is the affinity scoring gate (Affinity_Prob *≥* 0.5), which removes 79.9% of the pool in one step (1319 *→* 265, Figure 6). Molecular weight and LogP together remove a further 9.8% (265 *→* 239). The next four checks, hydrogen bond donor count, hydrogen bond acceptor count, topological polar surface area, and rotatable bond count remove no additional candidates, and synthetic accessibility removes only 1. The QED and novelty checks remove 11 and 9 further candidates respectively, leaving a final pool of 218, 16.5% of the original 1319. This combined cascade enforces the Affinity Probability threshold uniformly across pipelines, the per pipeline “filtered pool” figures used elsewhere in this report (Tables 1, 3 and 4) are drawn from each pipeline’s own SA,QED and property filtering and do not themselves enforce that affinity gate, which is why the two views of the data do not reconcile numerically (Figure 6).

### **B.3** Scoring methodology

Affinity, potency, and structural confidence values for every candidate come from a single cofolding based predictor, applied uniformly regardless of which pipeline generated the molecule. For each candidate the predictor returns a binary affinity probability, a regressed affinity value converted to the customary potency scale via pIC50 (and hence IC50 = 10*^−^*^pIC50^ *×* 10^6^ *µ*M), and structural confidence metrics reported here as Confidence (derived from predicted and interface template modelling scores, pTM/ipTM) and pLDDT, the per complex confidence in the modelled structure.

QED and SAS are computed independently by standard cheminformatics implementations and are not outputs of the affinity predictor.

### **B.4** Benchmarking Results

#### B.4.1 Benchmark Overview

The outcome of the benchmark is reported in Table 6. As in comparable benchmarks of generative chemistry pipelines, no single tool dominates the panel, and the ranking depends on the criterion adopted.

**Table 6:** Benchmark of five generative pipelines for ACVR1. Bold face marks the most favourable value of each metric; a dagger marks the least favourable.

| Pipeline | $N$ | Affinity probability | | | QED<sup>a</sup> | SAS<sup>b</sup> | Scaff. div. (%) | Intra-Tan.<sup>c</sup> |
| --- | --- | --- | --- | --- | --- | --- | --- | --- |
|  |  | Mean | Best | Top-5 |  |  |  |  |
| Pocket2Mol | 97 | 0.159 | 0.539 | 0.523 | 0.555 | **2.105** | 49.5 | **0.126** |
| SynFlowNet | 414 | 0.350 | 0.898 | 0.877 | 0.624 | 2.295 | 88.9 | 0.149 |
| REINVENT | 95 | 0.201 | 0.914 | 0.881 | 0.610 | 2.317 | 91.6 | 0.142 |
| LLMsFold | 172 | **0.815** | **0.957** | **0.955** | 0.585 | 2.517 | 24.4 | 0.540 |
| S3-GFN | 91 | 0.335 | 0.680 | 0.620 | **0.929** | 2.513 | **100.0** | 0.171 |

<sup>a</sup> Mean quantitative estimate of drug-likeness over each pipeline’s own top 5 candidates by affinity; higher is better.
<sup>b</sup> Mean synthetic accessibility score over the same top 5; lower is better.
<sup>c</sup> Mean pairwise Tanimoto similarity within each pipeline’s filtered pool (Morgan fingerprints, $r = 2$ , 2048 bit); lower values indicate greater internal chemical diversity.

### **B.5** Predicted affinity

LLMsFold (0.815) is unambiguously the strongest binder by pool wide mean affinity, Pocket2Mol (0.159) remains the panel’s weakest on this measure. The middle of the panel has been reshuffled by SynFlowNet’s mean affinity 0.350 is barely ahead of S3-GFN’s 0.335. SynFlowNet and REINVENT (0.201) remain competitive with LLMsFold on best (0.898 and 0.914) and top 5 mean (0.877 and 0.881) affinity.(Table 6).

### **B.6** Drug likeness and synthetic accessibility

S3-GFN redraws the boundary on drug likeness entirely by its top 5 QED (0.929) is not merely the best in the panel but sits far outside the range spanned by the other four pipelines (0.555–0.624). Its top-5 SAS (2.513) is now essentially tied with LLMsFold’s (2.517). Among the other four, no pipeline leads on every chemistry metric, SynFlowNet’s top candidates are the most drug like after S3-GFN (QED = 0.624), Pocket2Mol’s are the easiest to synthesise (SAS = 2.105), which is consistent with its small, fragment like output rather than deliberate optimisation for accessibility.

### **B.6.1** Full Pool Distributions

The top-5 statistics summarize each pipeline’s best candidates, which are not always representative of typical output. Per-molecule box plots of the full filtered pool (Figure 8, summarized in Table 7) show that Pocket2Mol illustrates this gap: its top-5 SAS (2.105) is the most favorable in the panel, yet its full-pool median SAS (*≈* 4.08) is the least favorable, with a right-skewed spread. Its easiest-to-make candidates represent an unrepresentative tail of an otherwise harder-to-synthesize pool. The same pattern holds for drug-likeness: full-pool median QED (*≈* 0.34) sits far below its top 5 mean (0.555).

**Table 7:** Full pool distribution summary (median [interquartile range]), read from per-molecule box plots of the filtered pool. Bold face marks the most favourable median in each column.

| Pipeline | Affinity_Prob | QED | SAS |
| --- | --- | --- | --- |
| Pocket2Mol | 0.15 [0.10–0.24] | 0.34 [0.19–0.58] | 4.08 [2.65–4.80] |
| SynFlowNet | 0.33 [0.26–0.41]<sup>b</sup> | 0.44 [0.34–0.55] | 3.08 [2.82–3.40] |
| REINVENT | 0.09 [0.05–0.25]<sup>a</sup> | 0.57 [0.42–0.69] | **2.53** [2.30–2.98] |
| LLMsFold | **0.81** [0.77–0.93] | 0.57 [0.53–0.61] | 2.97 [2.58–3.30] |
| S3-GFN | 0.31 [0.21–0.44] | **0.93** [0.90–0.95] | 2.62 [2.28–3.05] |

<sup>a</sup> REINVENT’s whisker extends up to $\approx 0.91$ via a distinct cluster of high affinity outliers, well above its own interquartile range.
<sup>b</sup> SynFlowNet’s whisker extends up to $\approx 0.90$ via the same kind of high affinity outlier cluster.

REINVENT’s affinity distribution is bimodal: the bulk of its pool clusters at a low median (*≈* 0.09), with a distinct group of high-affinity outliers extending up to *≈* 0.91 that pulls its top 5 statistics far above typical candidates. S3-GFN’s QED dominance is consistent across the pool: its full-pool interquartile range (*≈* 0.90–0.95) sits entirely above every other pipeline’s third quartile, confirming that high drug-likeness is standard for this method.

### **B.6.2** Computational Cost

Wall clock generation time was measured for the five pipelines and differs by just over three orders of magnitude across them (Table 8). REINVENT generated 985 raw candidates in 5 minutes (*≈*197 molecules per minute), Pocket2Mol needed 43 minutes for 235 (*≈*5.5 molecules per minute), LLMsFold needed 46 minutes to produce its full pool of 172 (*≈*3.7 molecules per minute), and SynFlowNet needed 300 minutes (5 hours) to generate 698 unique raw candidates (*≈*2.3 molecules per minute). Finally S3-GFN took 600 minutes to generate 91 unique raw candidates with *≈*0.15 molecules per minute was the slowest of all. The comparison across pipelines is not perfectly like for like and

**Table 8:** Observed wall clock generation time, this deployment.

| Pipeline | Runtime (min) | Molec.<sup>a</sup> | Throughput (mol min<sup>-1</sup>) |
| --- | --- | --- | --- |
| REINVENT | 5.0 | 985 | **197.0** |
| Pocket2Mol | 43.0 | 235 | 5.5 |
| LLMsFold | 46.2 | 172<sup>b</sup> | 3.7 |
| SynFlowNet | 300.0 | 698 | 2.3 |
| S3-GFN | 600.0 | 91 | 0.15 |

<sup>a</sup> REINVENT, Pocket2Mol, and SynFlowNet counts are raw generator output, prior to the cheminformatics cascade, not directly comparable to the filtered pool $N$ in Table 6 without accounting for that attrition.
<sup>b</sup> LLMsFold applies its filter cascade inline during generation, so this count is already the post filter pool used in Table 6.

### **B.7** Scaffold diversity and novelty

S3-GFN pushes scaffold diversity to its logical limit and all 91 of its filtered molecules resolve to 91 distinct Bemis Murcko scaffolds (100%), meaning no two candidates share even a core ring system. This is much higher than REINVENT scafold diversity already high 91.6% (87 of 95 molecules). SynFlowNet follows the same broad exploration pattern almost as strongly by 368 of its 414 filtered molecules resolve to distinct scaffolds (88.9%).

S3-GFN’s intra pipeline Tanimoto similarity (0.171) remains the highest of the four non LLMsFold pipelines, above SynFlowNet’s (0.149), REINVENT’s (0.142), and Pocket2Mol’s (0.126), indicating that while its molecules rarely share a ring system, they do draw repeatedly on similar substituents and linker chemistry (Figure 9). LLMsFold sits at the opposite extreme from S3-GFN and SynFlowNet on scaffolds (24.4% unique). The highest inter pipeline pair is LLMsFold and REINVENT (0.180), reflecting the partial overlap in aminopyridine biaryl chemotype visible in the UMAP projection (Figure 9).

**Figure 9:**
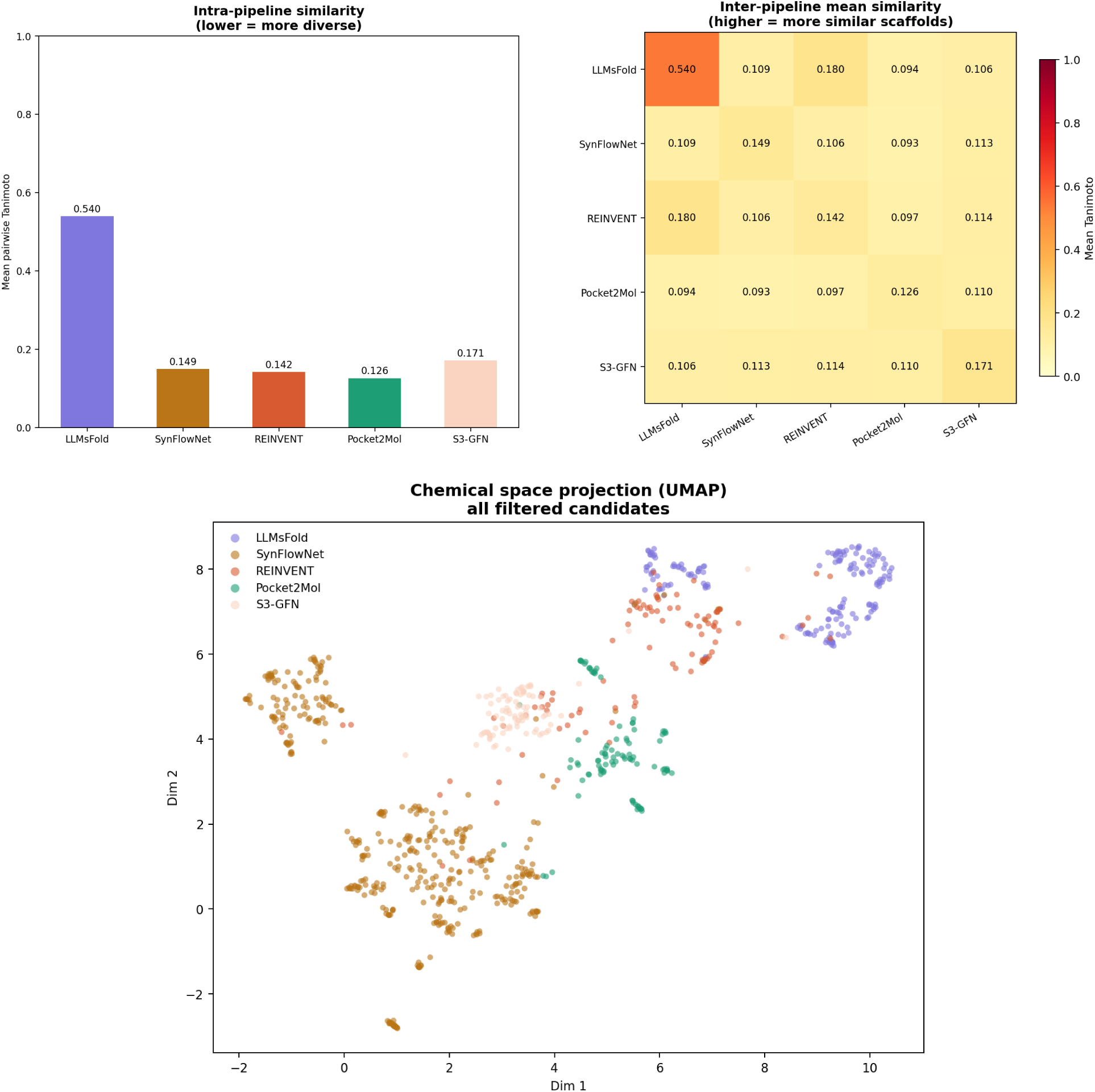
Tanimoto scaffold diversity analysis (Morgan fingerprints, *r* = 2, 2048 bit), five pipelines. Intra pipeline similarity (top left) is highest for LLMsFold (0.540) and lowest for Pocket2Mol (0.126). The inter pipeline heatmap (top right) shows the LLMsFold and REINVENT is the most similar pair (0.180). The UMAP projection (bottom) shows SynFlowNet occupying a large, distinct region of chemical space of its own, separate from the LLMsFold cluster, while S3-GFN continues to occupy its own separate region, bordering but largely apart from Pocket2Mol and REINVENT

**Table 9:** Bemis Murcko scaffold summary, property filtered pool (pre novelty gate).

| Pipeline | Filt. pool | Uniq. scaff. | Div. (%) |
| --- | --- | --- | --- |
| LLMsFold | 172 | 42 | 24.4 |
| SynFlowNet | 414 | 368 | 88.9 |
| REINVENT | 95 | 87 | 91.6 |
| Pocket2Mol | 97 | 48 | 49.5 |
| S3-GFN | 91 | 91 | 100.0 |

**Table 10:** Global composite score leaderboard: top 10 candidates, plus the only two entries outside LLMsFold to reach the global top 50 (SynFlowNet, ranks 49 and 50). All are flagged novel.

| Rank | Pipeline | Comp. | Affin. | pIC50 | SAS/QED |
| --- | --- | --- | --- | --- | --- |
| 1 | LLMsFold | 0.847 | 0.953 | 10.72 | 2.64 / 0.59 |
| 2 | LLMsFold | 0.843 | 0.949 | 10.51 | 2.81 / 0.64 |
| 3 | LLMsFold | 0.836 | 0.953 | 10.29 | 2.35 / 0.61 |
| 4 | LLMsFold | 0.836 | 0.953 | 10.74 | 2.58 / 0.55 |
| 5 | LLMsFold | 0.835 | 0.945 | 10.55 | 2.51 / 0.59 |
| 6 | LLMsFold | 0.831 | 0.949 | 10.81 | 2.45 / 0.51 |
| 7 | LLMsFold | 0.826 | 0.945 | 10.21 | 2.39 / 0.61 |
| 8 | LLMsFold | 0.825 | 0.953 | 10.82 | 2.43 / 0.49 |
| 9 | LLMsFold | 0.823 | 0.957 | 10.25 | 2.88 / 0.61 |
| 10 | LLMsFold | 0.820 | 0.949 | 10.48 | 2.92 / 0.58 |
| 49 | SynFlowNet | 0.751 | 0.879 | 9.61 | 2.17 / 0.65 |
| 50 | SynFlowNet | 0.742 | 0.859 | 9.43 | 2.12 / 0.71 |

### **B.8** Global composite ranking

Every novel candidate surviving the cascade in Figure 6 receives a composite score blending predicted affinity, drug likeness, and synthetic accessibility. This revision recovers the declared component weights: Affinity_Prob (0.40), pIC50 (0.25), QED (0.15), a synthesisability term SAS_inv (0.10), and structural Confidence (0.10), summing to This resolves part of the mystery raised in earlier revisions of this report, where a least squares fit using only Affinity_Prob, QED, and (10 *−* SAS)*/*9 against the top 50 candidates correlated reasonably well with a roughly affinity led blend (*r ≈* 0.90) but never reproduced the score exactly (*R*^2^ *≈* 0.88 at best). No REINVENT, Pocket2Mol, or S3-GFN candidate reaches the top 50 at all still falls well short of the *∼*0.94–0.96 affinity range that dominates this leaderboard.

### **B.9** Intellectual-property overlap

Novelty screening is not a formality, it changes which candidates lead each pipeline. Ranked by raw affinity before the novelty gate, LLMsFold’s own best hit (affinity = 0.957) is flagged as patented, SynFlowNet’s best hit (0.898) is a known compound, and REINVENT’s top two hits (0.914 and 0.898) are patented and known, respectively. Pocket2Mol keeps all but its weakest raw hit novel, and S3-GFN is the cleanest pipeline of all on this measure and every one of its top five raw hits is flagged novel. LLMsFold’s and SynFlowNet’s joint dominance of the leaderboard is explained by their combination of strong raw affinity and, unlike REINVENT, a deep enough bench of novel analogues once IP encumbered hits are removed and REINVENT’s two best raw hits are patented and known respectively, which knocks its most competitive candidates out of contention before the composite ranking is even computed.

## **C** Discussion

LLMsFold’s key advantage in this benchmark is that its strong performance requires no target specific adaptation. Unlike SynFlowNet, REINVENT, and S3-GFN, which all rely on GFlowNet or RL based post training against a reward signal defined for this specific pocket, LLMsFold is deployed directly, without any retraining or fine tuning step for ACVR1 and yet it still dominates the panel on the metric that matters most for a real campaign, predicted affinity. It leads on every affinity measure (mean, best, and top 5), and unlike SynFlowNet or REINVENT, this strength is not an artifact of a handful of standout candidates, its full 172 molecule pool sustains a mean Affinity Probability of 0.815, roughly 2.3*×* S3-GFN’s and more than 4*×* Pocket2Mol’s, so a randomly chosen LLMsFold candidate is still a strong binder, not just its best five. This translates directly into the composite leaderboard, where LLMsFold supplies 48 of the top 50 candidates essentially unchallenged. Its drug likeness and synthetic accessibility, while not best in class, are solidly mid panel and far from the extremes seen in Pocket2Mol’s fragment like output, giving it a broadly balanced profile rather than a narrow one metric win. Taken together, LLMsFold offers the combination that matters for near term lead generation, no per target training overhead, the highest and most consistent affinity in the panel, and drug like chemistry that doesn’t need to be traded off to get there a combination none of the reaction constrained or RL tuned baselines, including S3-GFN, currently match.

## Footnotes

4 https://groq.com/

5 https://developer.nvidia.com/dgx-cloud/nvcf

6 Version 1.11.1, released on January 23, 2026

7 https://github.com/MolecularAI/REINVENT4

